# Population-level HIV incidence estimates using a combination of synthetic cohort and recency biomarker approaches in KwaZulu-Natal, South Africa

**DOI:** 10.1101/306464

**Authors:** Eduard Grebe, Alex Welte, Leigh F. Johnson, Gilles van Cutsem, Adrian Puren, Tom Ellman, Jean-François Etard, the Consortium for the Evaluation and Performance of HIV Incidence Assays (CEPHIA), Helena Huerga

## Abstract

**Introduction:** There is a notable absence of consensus on how to generate estimates of population-level incidence. Incidence is a considerably more sensitive and harder to estimate indicator of epidemiological trends than prevalence. We used a novel hybrid method to estimate HIV incidence by age and sex in a rural district of KwaZulu-Natal, South Africa.

**Methods:** Our novel method uses an ‘optimal weighting’ of estimates based on an implementation of a particular ‘synthetic cohort’ approach (interpreting the age/time structure of prevalence, in conjunction with estimates of excess mortality) and biomarkers of ‘recent infection’ (combining Lag-Avidity, Bio-Rad Avidity and viral load results to define recent infection, and adapting the method for age-specific incidence estimation). Data were obtained from a population-based cross-sectional HIV survey conducted in Mbongolwane and Eshowe health service areas in 2013.

**Results:** Using the combined method, we find that age-specific HIV incidence in females rose rapidly during adolescence, from 1.33 cases/100 person-years (95% CI:0.98,1.67) at age 15 to a peak of 5.01/100PY (4.14,5.87) at age 23. In males, incidence was lower, 0.34/100PY (0.00-0.74) at age 15, and rose later, peaking at 3.86/100PY(2.52-5.20) at age 30. Susceptible population-weighted average incidence in females aged 15-29 was estimated at 3.84/100PY (3.36-4.40), in males aged 15-29 at 1.28/100PY(0.68-1.50) and in all individuals aged 15-29 at 2.55/100PY (2.09-2.76). Using the conventional recency biomarker approach, we estimated HIV incidence among females aged 15-29 at 2.99/100PY (1.79-4.36), among males aged 15-29 at 0.87/100PY(0.22-1.60) and among all individuals aged 15-59 at 1.66/100PY (1.13-2.27).

**Discussion:** HIV incidence was very high in women aged 15-30, peaking in the early 20s. Men had lower incidence, which peaked at age 30. The estimates obtained from the hybrid method are more informative than those produced by conventional analysis of biomarker data, and represents a more optimal use of available data than either the age-continuous biomarker or synthetic cohort methods alone. The method is mainly useful at younger ages, where excess mortality is low and uncertainty in the synthetic cohort estimates is reasonably small.

**Conclusion:** Application of this method to large-scale population-based HIV prevalence surveys is likely to result in improved incidence surveillance over methods currently in wide use. Reasonably accurate and precise age-specific estimates of incidence are important to target better prevention, diagnosis and care strategies.

## Introduction

HIV epidemic surveillance largely relies on cross-sectional measurements of prevalence, often by means of representative household surveys. However, for a non-remissible condition with extended survival time like HIV, instantaneous prevalence reflects the epidemic trajectory (incidence, mortality and migration) over a significant period prior to the survey. Estimating HIV incidence – the most sensitive and informative indicator of current epidemiological trends – therefore poses significant methodological challenges.

The ‘gold standard’ method of directly observing new infections in cohorts of HIV-negative individuals followed up over time are costly and logistically challenging, and it is difficult to ensure sufficient population representivity to ensure results can be generalised. Several alternative approaches have been proposed for estimating HIV incidence, including a ‘synthetic cohort’ approach – i.e. inferring incidence from the age and/or time structure of prevalence [1–6], from biomarkers for ‘recent infection’ measured in cross-sectional surveys [7–11], or using dynamical population models that have been calibrated to survey data [12–15]. No single method by itself achieves the desired levels of accuracy and precision [16].

In this work we develop a novel hybrid method which uses an ‘optimal’ weighting of, (a) an implementation of the ‘synthetic cohort’ approach of Mahiane et al. [6] – i.e. interpreting the age and time structure of prevalence, in conjunction with excess mortality – and (b) an adaptation of the Kassanjee et al. estimator for incidence from biomarkers of recent infection [8] that takes account of the age structure of recent infection (amongst HIV-positive individuals). The method of Mahiane et al. relies on the instantaneously exact, fully age-and time-structured, representation of the dynamical relation of prevalence, excess mortality and incidence. In the case of a relatively stable epidemic (i.e. relatively slow change in age-specific prevalence over time), the age structure of prevalence provides fairly precise age-specific incidence estimates.

We applied this method to a cross-sectional household survey conducted in a district of KwaZulu-Natal province (KZN) to estimate the HIV incidence by age and sex in the area at the time of the survey (2013). For the present analysis we assume stability, but we investigate the impact of plausible time-gradients of prevalence in the sensitivity analysis. Precision of incidences estimates is markedly lower for ages over 30, and we therefore report as primary results incidence over age range 15-29 years.

## Methods

### Survey design and procedures

The data analysed in this study were obtained from the Mbongolwane and Eshowe HIV Impact in Population Survey, conducted in 2013 in Mbongolwane, a rural area, and Eshowe, the main town in the uMlalazi Municipality in KZN, South Africa. A two-stage stratified clustered sampling strategy was used for the selection of households according to the 2011 Census, which indicated a population of approximately 120,000 at the time of the survey [17]. Individuals aged 15-59 years old living in sampled households, and who provided informed consent, were enrolled in the study.

The University of Cape Town Human Research Ethics Committee (HREC 461/2012), the Health Research Committee of the Health Research and Knowledge Management Unit of the KwaZulu-Natal Department of Health and the Comité de Protection de Personnes de Paris in France approved the study protocol.

Face-to-face interviewer-administered questionnaires were used to collect information on socio-demographics and sexual history at the sampled household. HIV testing, including pre-and post-test counselling, was done by certified lay counsellors, on site, using the Determine Rapid HIV-1/2 Antibody test kit as a screening test followed, in the case of a positive result, by the Unigold Rapid HIV test kit for confirmation. Venous blood specimens were collected from all participants who consented. HIV antibody-positivity was determined using the on-site rapid result, confirmed by laboratory-based ELISA in the case of discordant rapid test results. Specimens from participants confirmed to be HIV antibody-positive were subjected to the Sedia Limiting Antigen Avidity EIA (LAg) assay [18], the Bio-Rad Avidity assay [19], as well as a quantitative viral load, CD4 count and an ARV presence test. Viral load testing was performed using a NucliSens EasyQ HIV-1 v2.0 assay from Biomerieux. Qualitative testing for ARV drug levels, including nevirapine, efavirenz and lopinavir, was performed using a LC MS/MS qualitative assay. In addition, to detect acute infections in antibody-negative participants, HIV-negative specimens were subjected to pooled Nucleic Acid Amplification Testing (NAAT) testing (in 5-member pools) using Roche AMPLISCREEN, and specimens from positive pools tested individually using the Roche CAP/CTM assay.

More detail on the survey has been published elsewhere [17,20–22].

### Estimating incidence using biomarkers for ‘recent’ infection

We used calibration data from the Consortium for the Evaluation and Performance of HIV Incidence Assays (CEPHIA) to explore a range of recent infection case definitions based on combinations of LAg normalised optical density (ODn), Bio-Rad Avidity index (AI) and viral load thresholds, and selected an ‘optimal’ recent infection testing algorithm (RITA) based on the variance of the incidence estimates produced. The procedures and results are detailed in Appendix 1: Optimal RITA identification and calibration. These show that a RITA that defines recent infection as NAT+/Ab− OR Ab+/ODn < 2.5/AI < 30/VL > 75 achieves a mean duration of recent infection (MDRI, adjusted for the sensitivity of the screening algorithm) of 217 days (95% CI: 192,244) and a context-specific false-recent rate (FRR) of 0.17% (95% CI: 0.05%,0.35%). We analyse sensitivity to imperfectly-estimated FRR in Appendix 2: Sensitivity analyses.

This definition of recency was then employed to estimate incidence in the study population, by age group and sex, using the method of Kassanjee et al. [8]. The well-known Kassanjee et al. estimator, adapted for use in complex surveys by allowing the use of proportions and their standard errors, rather than survey counts, is given in Eq 1.
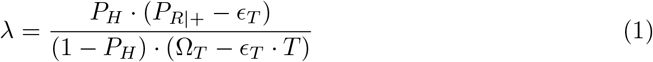

where *P*_*H*_ is the prevalence of HIV, *P*_*R*|+_ is the proportion of recency tests performed on HIV-positive participants that produced a ‘recent’ result, is the MDRI and Ω_*T*_ is the FRR, and *ϵ*_*T*_ is the chosen time cutoff beyond which a ‘recent’ result is considered ‘falsely recent’ by definition. Note that the product of *P*_*H*_ and *P*_*R*|+_ is the overall prevalence of recency in the sample. This estimator is implemented in the inctools R package [23]. The documentation of inctools provides details on estimating the variance of incidence estimates using both delta method and bootstrapping approaches.

Owing to the small ‘recent infection’ case counts, statistical uncertainty reaches unacceptable levels when age groups are small, and we therefore estimated incidence using the conventional approach in 15 to 29 year-olds and 30 to 59 year-olds.

For the purpose of the combined method described below, we further adapted the estimator for age-dependent prevalence of recent infection, allowing us to estimate highly granular age-specific incidence using the recent infection biomarker data in the survey. Details are provided in the section on the combined method.

### Estimating age-specific incidence using the Mahiane et al. ‘synthetic cohort’ method

We employ the incidence estimator of Mahiane et al. [6] to estimate incidence from the age structure of prevalence. The estimator was derived from the fundamental relationship between incidence, prevalence and mortality in a non-transient condition – with prevalence viewed as the accumulated incidence over time, accounting for the removal of prevalent cases from the population through condition-induced ‘excess’ mortality. This is shown using a simple dynamical SI-type model, where it is demonstrated that simply rearranging the differential equations describing change in the state variables for the susceptible and infected groups yields an estimator for incidence that relies only on prevalence and excess mortality (but critically, not total mortality). In an age-structured population, this approach yields the incidence estimator in Eq 16.
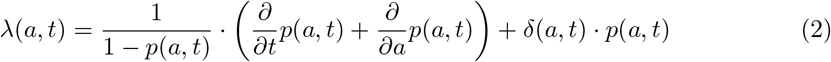

where *p*(*a*,*t*) is age and time-specific prevalence and *δ*(*a*,*t*) is age and time-specific excess mortality.

In a stable epidemic, where the age structure of prevalence is not changing at a significant rate in secular time (see discussion section), the age-structure of prevalence from a single cross-sectional prevalence survey is informative, and the estimator can be simplified to:

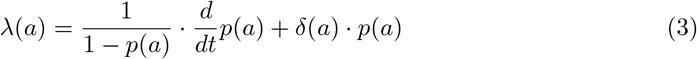

We obtained age-specific incidence estimates by fitting a regression model for prevalence as a function of age to finely-grained data (i.e., not using integer ages, but the difference in days between the birth date and interview date of each participant), using a generalised linear model with a cubic polynomial in age as predictors and a logit link:

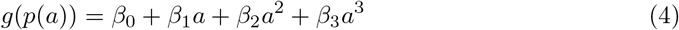

with *g*() the logit link function, so that

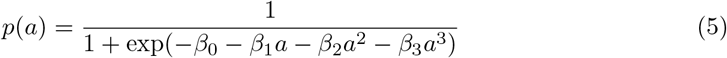

and

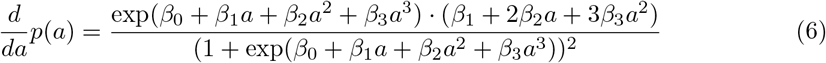

We fit the model, separately for males and females, to data from participants aged 15 to 34 years. This provided us with a continuous function, *p*(*a*), for 15 ≤ *a* < 35. We derived, for each sex, the function for excess mortality, *δ*(*a*), from age-specific AIDS mortality estimates for KwaZulu-Natal province produced by the Thembisa demographic model [24], allowing us to estimate age-specific incidence, λ (*a*).

Reproducibility of the incidence estimate at any given age was investigated by bootstrapping the dataset (reproducing the complex sampling frame employed in the survey), refitting the models and obtaining an incidence estimate for each of the 10,000 resampled datasets. The standard deviation of the obtained estimates was computed to approximate the standard error, and the 2.5th and 97.5th percentiles to approximate the 95% confidence interval.

### Estimating age-specific incidence using the combined method

In order to estimate age-specific incidence by combining HIV prevalence data and biomarkers for recent infection, available in the same dataset, we estimated age-specific incidence (and its variance) using (1) the synthetic cohort method described above, and (2) an adaptation of the Kassanjee et al. estimator to age-structured recency biomarker data. The adapted estimator is shown in Eq 7.
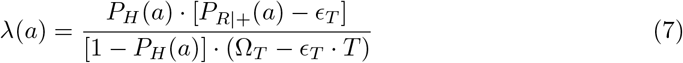

where *P*_*H*_(*a*) is the age-specific prevalence of HIV (estimated as in the previous section), and *P*_*R*|+_ (*a*) is the age-specific prevalence of recency amongst HIV-positives (described below).

In order to obtain the prevalence of recency as a function of age we fit a generalised linear regression model with log of age as linear predictor and a complementary log-log link. This functional form implies an exponential decline in the prevalence of recency with age, which captures the epidemiologically sensible assumption that at young ages larger proportions of infections were acquired in the recent past. The model has the functional form:

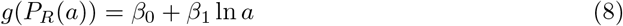

with *g*() the link function, so that

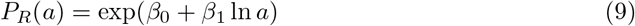

Owing to the use of prevalence in both estimates, the incidence estimates are necessarily correlated. We therefore resample the data (replicating the complex sampling frame), fit the models of *P*_*H*_(*a*) and *P*_*R*_(*a*), and at each age of interest, obtain the two incidence estimates, λ_*P*_ (incidence from age-structured prevalence) and λ_*R*_ (incidence from age-structured recency amongst positives). We then evaluate, at each age of interest, from 10,000 bootstrap iterations, the variances and covariance of the two incidence estimates, in the case of λ_*R*_ further incorporating uncertainty in MDRI and FRR. We then compute a combined incidence estimate using a weighted average of the two estimates. The implied weighting function, *W*(*a*), derived from the ‘optimal’ weighting factors, *W*_*a*_ (i.e. the weighting factor that minimises variance of the combined estimate) obtained at each evaluated age, is then convolved with the combined incidence function. At a particular age *a*, incidence from the combined method is given by Eq 10.
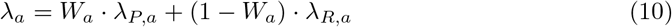

with 0 ≤ *W*_*a*_ ≤ 1, and consequently no normalisation to total weight is required. The variance of the incidence estimate at a given age is obtained from Eq 11.
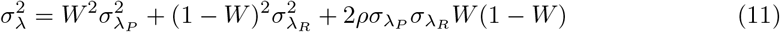

with *ρ* the Pearson‘s correlation coefficient between λ_*P*_ and λ_*R*_ at that age. The value of *W* that minimises total variance at the age of interest is obtained from the following formula, derived by setting Eq 11 to 0 and solving for *W*:

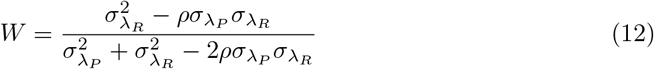

The continuous incidence function λ(*a*) is then obtained by fitting a cubic interpolating spline (using the method of Forsythe, Malcolm and Moler) to estimated incidence, λ_*a*_, for all ages in the range 15 to 35 years, evaluated at steps of 0.25 years.

For comparability with conventional age-group estimates, ‘average incidence’ was estimated in age bins. The integral of the λ(*a*) function was evaluated over the age range for which average incidence was sought, and weighted using a weighting function reflecting (a) the sampling density, or (b) the susceptible population density, to obtain average incidence. For population weighting, the population by age and sex was obtained from the 2011 Census for Eshowe and Mbongolwane, and the susceptible population size estimated using prevalence estimates from the survey data. Susceptible population-weigthted estimates are presented as primary results.

The unweighted incidence spline function, λ(*a*) was weighted by a weighting function *f*(*a*), derived from either the sampling density or the susceptible population density, and the integral evaluated over the age range of interest (*a*_0_ to *a*_1_) in order to obtain weighted average incidence over that range, as shown in Eq 13.
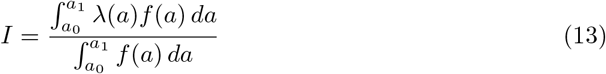

The procedure was performed separately for males and females, and in order to obtain overall average incidence, these estimates were then further weighted using the weighting functions for males and females.

Defining total weights for the two sexes as 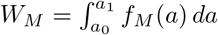 and 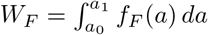, for any age interval and weighting function, the total incidence is then given by Eq 14.
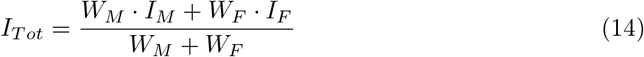

Confidence intervals were obtained by bootstrapping the data (10,000 iterations), and in each iteration estimating average incidence.

### Sensitivity analyses

In order to investigate the sensitivity of our analyses to uncertainty in the False-Recent Rate, we repeated the incidence estimation procedure using a range of FRRs between 0% and 1%. We further investigated sensitivity of average incidence to the weighting scheme.

The implementation of the method developed in this paper does not take into account change in prevalence (and incidence) in the time dimension. This is valid when the epidemic is relatively stable and most of the information is captured in the age structure of prevalence. In order to investigate sensitivity to possible change over time in age-specific prevalence, we investigated a number of hypothetical scenarios in which age-specific prevalence is increasing or decreasing exponentially.

Sensitivity analyses are reported in Appendix 2: Sensitivity analyses.

## Results

Conventional analysis of the biomarkers for recent infection (in large age bins) yielded an overall HIV incidence estimate for individuals aged 15 to 59 years at the time of the survey of 1.60 cases/100 person-years (PY) (95% CI: 1.11,2.16). In males 15-59 the incidence was estimated at 0.71/100PY (0.22,1.25) and in females 15-59 at 2.26/100PY (1.48,3.14). Among individuals aged 15-29, the main group of interest in this work, overall incidence was estimated at 2.03/100PY (1.37,2.77), for males at 0.89/100PY (0.28,1.58) and for females at 3.05/100PY (1.87,4.37). These results are presented in Table 5. Smaller age bins do not yield informative results using the conventional approach, owing the small case counts of recent infections.

**Table 1.** Conventional biomarker-based estimates for large age groups.

| Age group<br>*years* | Males<br>P.E. (95% CI)<br>*cases/100PY* | Females<br>P.E. (95% CI)<br>*cases/100PY* | Total<br>P.E. (95% CI)<br>*cases/100PY* |
| --- | --- | --- | --- |
| [15,30) | 0.89 (0.28,1.58) | 3.05 (1.87,4.37) | 2.03 (1.37,2.77) |
| [30,60) | 0.22 (0.00,0.99) | 1.06 (0.15,2.04) | 0.78 (0.12,1.48) |
| Overall [15-60) | 0.71 (0.22,1.25) | 2.26 (1.48,3.14) | 1.60 (1.11,2.16) |

Implicitly weighted by sampling density.

By way of comparison, the age-continuous biomarker-based method, which is a key component of the combined method, yielded ‘average incidence’ estimates (weighted by susceptible population density) in individuals aged 15-29 overall of 1.87 cases/100PY (1.31,2.43), in males of 0.81/100PY (0.22,1.45) and in females of 2.95/100PY (1.98,4.04). The synthetic cohort method yielded susceptible population-weighted average incidence in individuals aged 15-29 overall of 3.19/100PY (2.83,3.56), in males of 2.00/100PY (1.53,2.46) and in females of 4.39/100PY (4.00,4.85). Using the combined method, we obtained incidence estimates in individuals aged 15-29 of 2.54/100PY (2.07,2.77), in males of 1.26/100PY (0.64,1.49) and in females of 3.83/100PY (3.35,4.37). These results (as well as for five-year age bins) are reported in Table 6 and Table 3.

**Table 2.** ‘Average incidence’ estimates by age group using the biomarker and synthetic cohort methods.

| Age group<br>*years* | Age-continuous biomarker mothod |  |  | Synthetic cohort method |  |  |
| --- | --- | --- | --- | --- | --- | --- |
|  | Males | Females | Total | Males | Females | Total |
|  | P.E. (95% CI)<br>*cases/100PY* | P.E. (95% CI)<br>*cases/100PY* | P.E. (95% CI)<br>*cases/100PY* | P.E. (95% CI)<br>*cases/100PY* | P.E. (95% CI)<br>*cases/100PY* | P.E. (95% CI)<br>*cases/100PY* |
| [15,20) | 0.45 (0.08,0.88) | 1.92 (1.02,3.05) | 1.20 (0.73,1.78) | 0.27 (0.00,0.60) | 2.90 (2.30,3.48) | 1.61 (1.24,1.92) |
| [20,25) | 0.71 (0.16,1.38) | 3.73 (2.41,5.30) | 2.20 (1.51,2.92) | 1.59 (1.14,2.05) | 5.26 (4.53,6.20) | 3.40 (2.96,3.91) |
| [25,30) | 1.50 (0.40,2.73) | 3.62 (2.04,5.38) | 2.52 (1.52,3.45) | 5.09 (3.36,6.64) | 5.70 (4.33,7.30) | 5.38 (4.23,6.65) |
| [30,35) | 2.02 (0.24,4.50) | 2.66 (0.71,5.47) | 2.33 (0.99,4.30) | 4.07 (0.00,11.18) | 8.53 (2.29,16.43) | 6.20 (2.47,11.29) |
| [15,30) | 0.81 (0.22,1.45) | 2.95 (1.98,4.04) | 1.87 (1.31,2.43) | 2.00 (1.53,2.46) | 4.39 (4.00,4.85) | 3.19 (2.83,3.56) |

Weighted by susceptible population density.

**Table 3.**
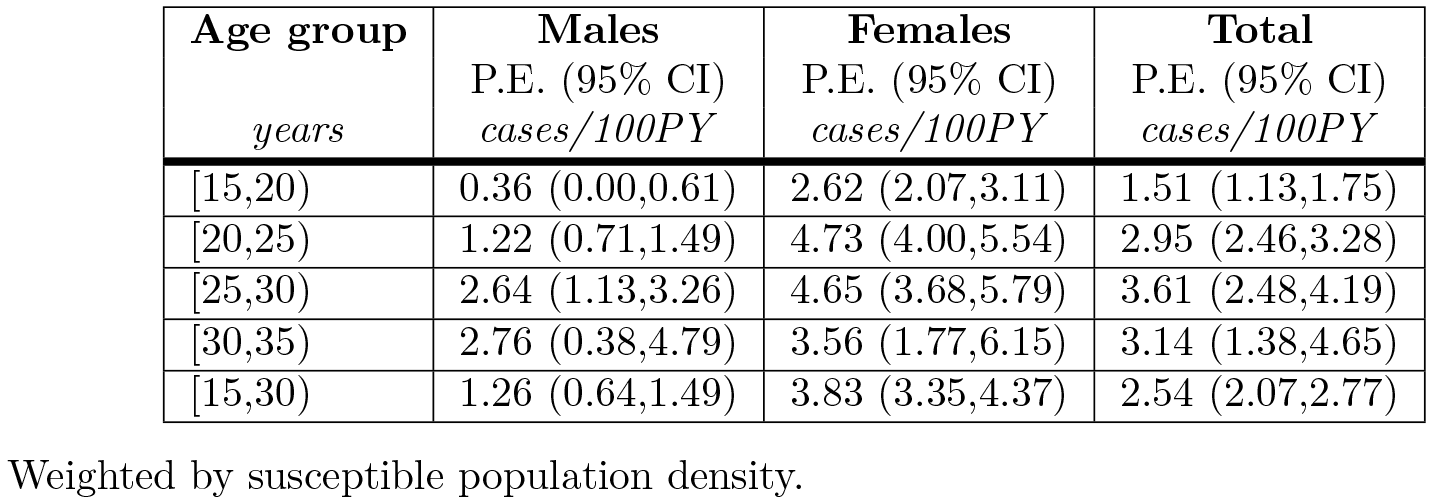
‘Average incidence’ estimates by age group using combined method.

Age-specific estimates using the combined method are shown in the figures. Incidence estimates are presented as continuous functions of age for individuals aged 15-29, with the contributions of the age-continuous biomarker and synthetic cohort methods. Fig 5 shows the overall results, Fig 6 the estimates for males and Fig 7 the estimates for females. Estimates become uninformative at ages over 30, owing to greatly increased statistical uncertainty.

Incidence in females rose steeply during the teenage years, from 1.31 cases/100PY (0.97,1.66) at age 15 to a peak of 4.95/100PY (4.09,5.81) at age 23. Incidence was lower but still very high – in women in their late twenties and early 30s, with estimated incidence of 4.50/100PY (3.07,5.92) at age 30. Uncertainty in the estimates increased with age (with a standard error of approximately 0.7 at age 30, compared to 0.4 at age 23). Estimates were very imprecise for ages over 30: at age 35, incidence was estimated at 2.78/100PY, with a standard error of 1.67, resulting in a 95% CI of 0.00,6.06. Age-specific incidence in teenaged males was substantially lower than in females, estimated at 0.32 cases/100PY (0.00,0.65) at age 15, and rising sharply from the early twenties, peaking at 4.10/100PY (2.75,5.46) at age 30. Incidence in males aged 23 was estimated at 1.39/100PY (0.95,1.82). Overall incidence estimates reflect the estimates for males and females so that estimated incidence at age 15 was 0.82 cases/100PY (0.64,1.00), peaked at 4.47/100PY (3.52,5.41) at age 29, and was 4.42/100PY (3.37,5.47) at age 30.

**Fig. 1.**
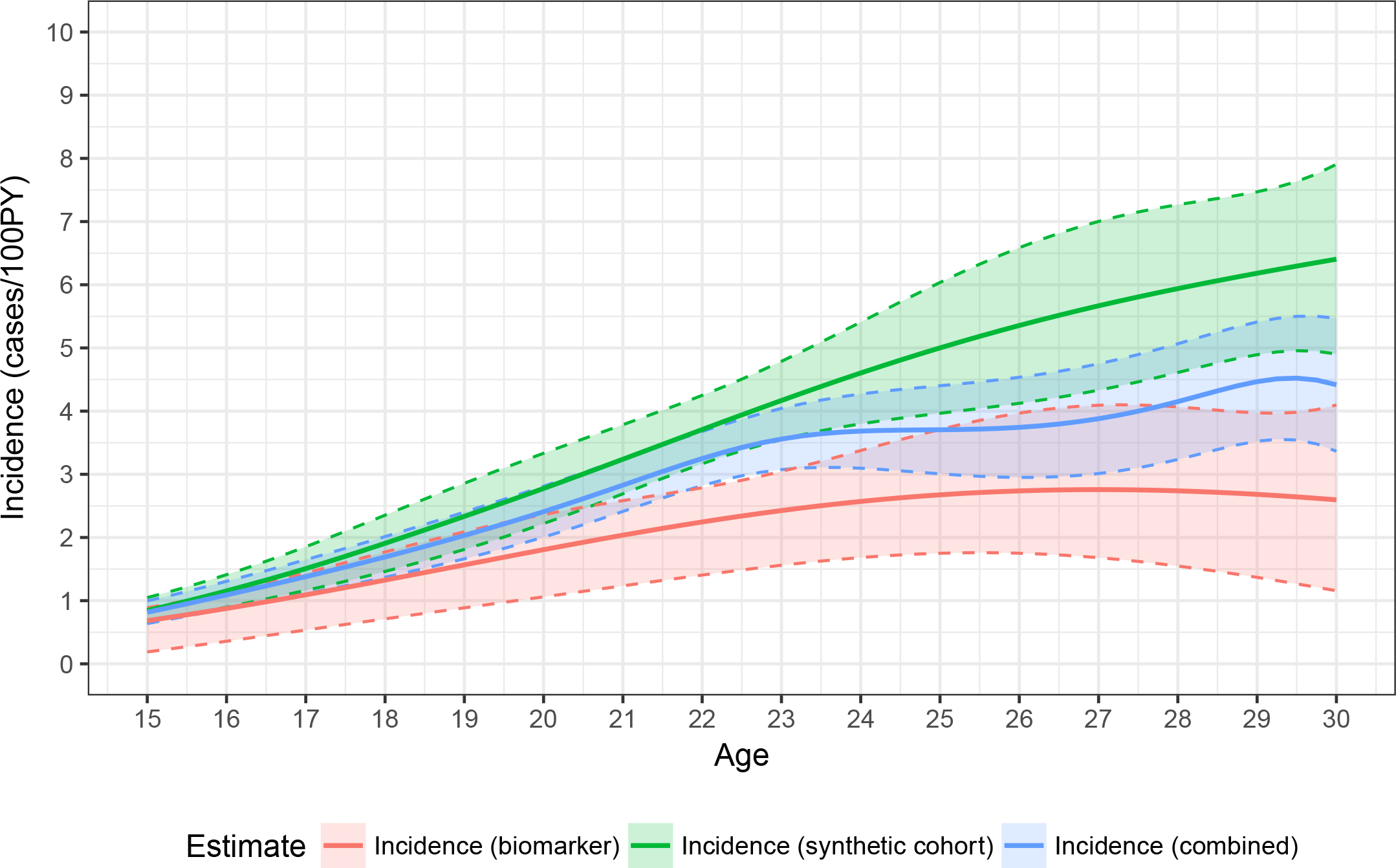
HIV incidence by age in males and females aged 15-30, using the synthetic cohort, recency biomarker and combined methods.

## Discussion

This study describes a novel hybrid method that allows for reasonably precise estimation of age-specific incidence up to about age 30 years. It constitutes a significant improvement over conventional cross-sectional incidence estimation using biomarkers of recent infection, where small case counts limit informative estimates to large age bins.

We confirm previously-described very high incidence among young women and also among slightly older young men. A compartmental mathematical model developed by Blaizot et al. [25] produced similar incidence estimates by sex and age group when calibrated to the same data [26]. In females, incidence peaked at age 23, and in males at age 30. We have previously described that young people were more likely to transmit HIV. In the same survey, among individuals aged 15-19 years and 20-34 years 34% and 35% respectively were unaware of their HIV status and 66% and 53% were virally unsuppressed; both factors were associated with higher-risk sexual behaviour [20]. Precise age-specific incidence estimates are important to identify the age and gender groups most at risk. These findings highlight the need for targeted prevention and HIV testing strategies for girls and young women, as well as men aged 20 to 40 years.

The conventional biomarker-based approach does not allow finely-grained age-specific incidence estimation, since small case counts (or sample proportions) result in very wide confidence intervals. Even analysis of the data in five-year age bins produce estimates that cannot be clearly distinguished from zero. Our adapted age-continuous biomarker estimates provide reasonably reproducible estimates in younger individuals, where the parameterisation of the prevalence of recent infection (amongst HIV-positive individuals) is likely to be sound. However, this method would be more challenging to implement in older individuals, where the distribution of recent infections is more complicated.

**Fig. 2.**
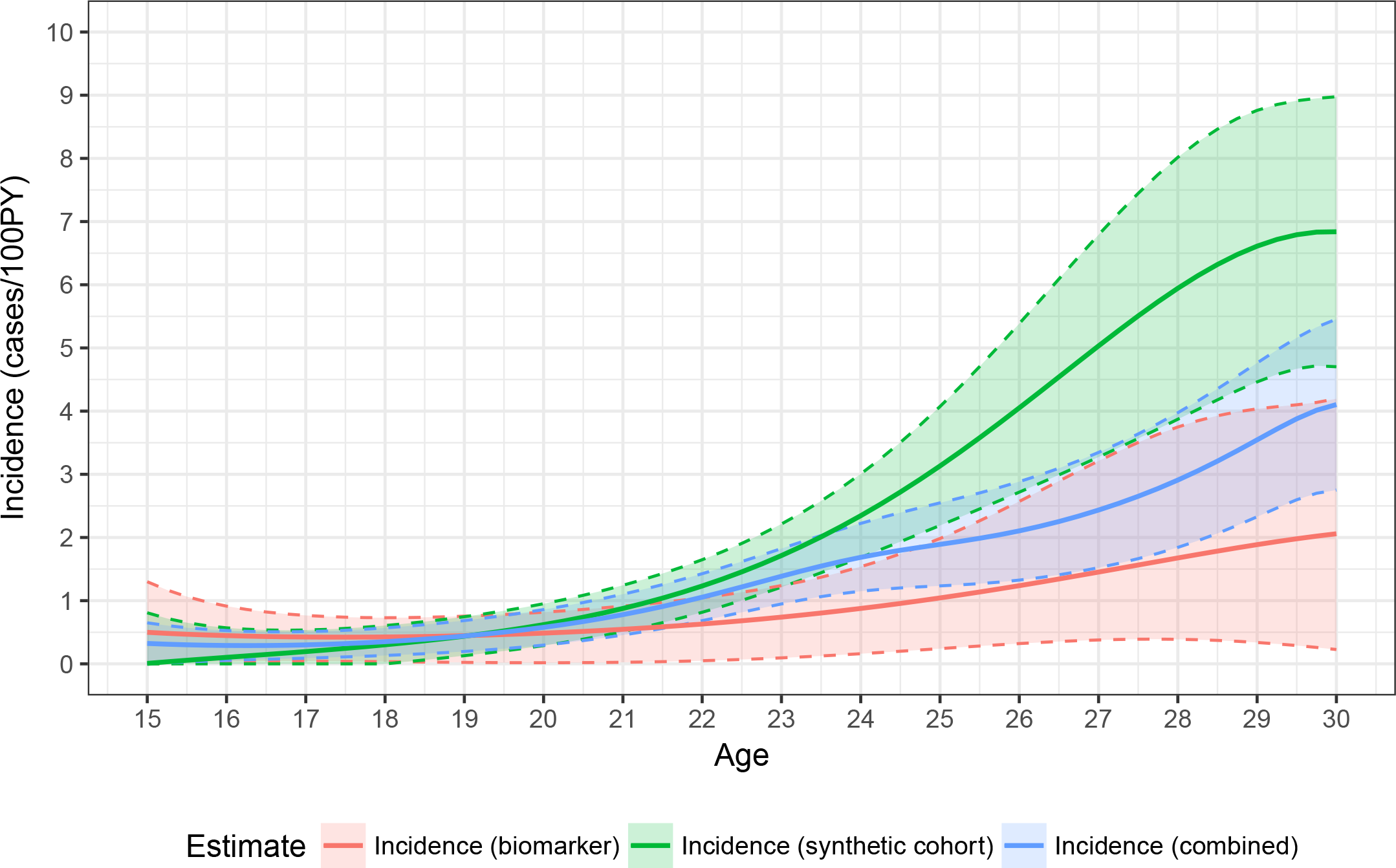
HIV incidence by age in males aged 15-30, using the synthetic cohort, recency biomarker and combined methods.

The synthetic cohort method provides additional information on incidence, and in certain age ranges is in fact more informative than the biomarker method. As can be seen in the figures, at younger ages the two estimates are very similar, but diverge at older ages. At younger ages the synthetic cohort method has greater precision (estimates have lower variance). In females, the combined estimate is weighted in favour of the synthetic cohort method throughout the age range 15 to 29, but with more heavily skewed at younger ages (weighting factor of 0.84 at age 15 and 0.54 at age 29), whereas in males the weighting tips towards the biomarker method at age 25.

The idea of using demographic structure of prevalence data to infer incidence is certainly not new. Williams et al. [1] developed something very close to the approach we are taking – the main difference being their proposal (in light of data available at that time) to use age-averaged rather than age-specific estimates of time dependence of prevalence. We follow the instantaneously exact, fully age and time-structured, representation of the relation of prevalence, mortality and incidence that was introduced in Mahiane et al. [6]. That paper also considered the previously-published methods of Brunet and Struchiner [2,3], Hallett et al. [4], and Brookmeyer and Konikoff [5], all of which were found to have substantial biases, noted to be the result of their various particular forms of dynamical approximation – essentially using assumptions of constant prevalence in age and time ranges.

**Fig. 3.**
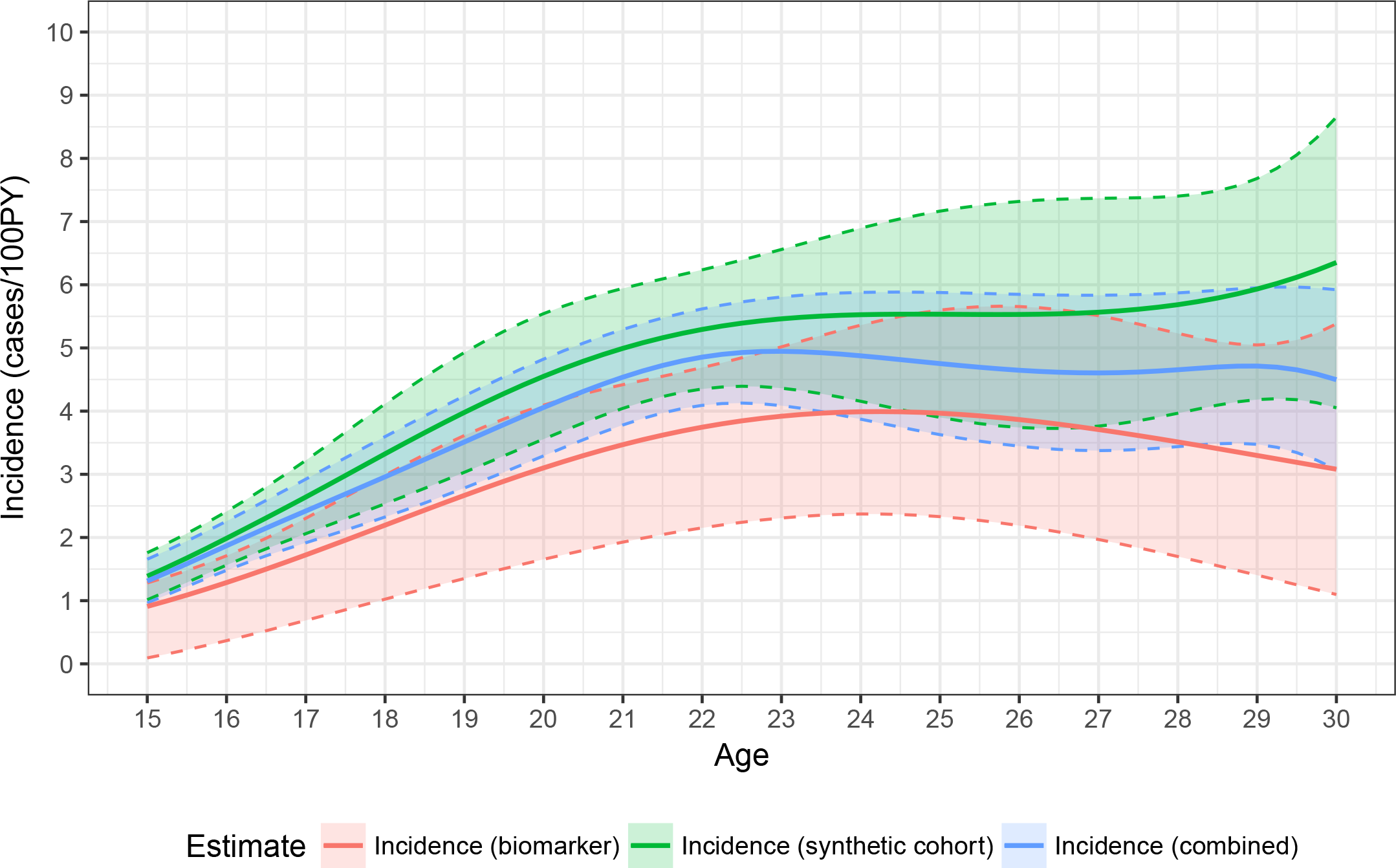
HIV incidence by age in females aged 15-30, using the synthetic cohort, recency biomarker and combined methods.

A major advantage of the hybrid approach is that it combines both age-specific HIV prevalence data and biomarker data, thus reducing the risk of bias in HIV incidence estimation. By combining the estimates from the two methods, age-specific incidence can be estimated with significantly greater precision than with the biomarker method alone. For example, at age 15 in females, the standard error on the age-continuous biomarker estimate of 0.91 cases/100PY is 0.42 (i.e. a coefficient of variation of 46%) and the standard error on the weighted average of 1.31/100PY is 0.18 (CoV = 13%). The narrower confidence bounds around the combined method estimates can be clearly seen in the figures. At certain ages, there is a very substantial improvement, for example in males aged 22, the CoV on the biomarker estimate is 47%, while on the combined estimate it is 18%. The precision of the combined method is not greatly enhanced over that of the synthetic cohort method, but estimates are likely to be more accurate, especially where information on change in the age structure of prevalence over time is not available, which may bias estimates.

For the recent infection case definition adopted for this analysis we estimated, using CEPHIA calibration data, a very small context-specific false-recent rate. Unfortunately, the FRR is also the test property that is hardest to estimate, and where the transferability from calibration data to the surveyed population is most problematic. While we adopted a sophisticated approach to context adaptation of the test property estimates, these challenges remain. For present purposes we assumed that the test properties (MDRI and FRR) do not vary with age, although it is likely that the longer (on average) time-since-infection in older individuals would impact the FRR and that biological changes in the immune system would impact the MDRI. In estimating context-specific FRR we make assumptions about the population-level distribution of times-since-infection, but a lack of data on past incidence precludes a more nuanced age-specific FRR estimate. This limitation is addressed by means of a sensitivity analysis with respect to FRR, as reported in Appendix 2: Sensitivity analyses. Given the very low population-level FRR estimate, it is unlikely that this assumption introduces substantial bias. The sensitivity analyses indicate that our results are not highly sensitive to the false-recent rate, although it becomes more so at older ages, where the combined method relies more on the biomarker-based estimate.

A major limitation of this study is that we are analysing data from a single cross-section survey, providing no information on change in the (age-structured) prevalence over time. A second survey is planned in the study population, which would allow future analyses to be conducted that explicitly incorporate change over time. We investigated the sensitivity of our estimates to prevalence changes in time, and found that estimates from the combined method are not very sensitive to plausible rates of change in prevalence at the time of the survey. However, if the assumption of a stable epidemic were violated and rapid increases or declines in prevalence were taking place at the time of the survey, our method would exhibit significant bias. It would therefore be preferable to explicitly incorporate the time dimension in the analysis (by using data from serial prevalence surveys) and it is essential that the version of the method that ignores time is only applied in settings where the assumption of stability is sound.

Further, estimates become very uncertain at ages over 30 (see Fig 4), resulting in synthetic cohort, biomarker and combined method estimates with confidence intervals that stretch from close to zero to very large values. The failure of the method to provide informative estimates at higher ages requires further investigation. This limitation may, in part, reflect the particular parameterisation of regression models for HIV prevalence and for the prevalence of ‘recent infection’ used in the present analysis.

## Conclusion

This analysis demonstrates the value of age-structured prevalence data, when reasonable estimates of excess mortality are available, and that when additional biomarkers of recent infection are available these can be sensibly incorporated into age-specific incidence estimates. The novel hybrid method used in this analysis can be extended to allow the analysis of serial prevalence (and, when available, recent infection) data, without significant further conceptual development, for maximally informative incidence estimation. Application of this method to large-scale population-based HIV prevalence surveys is likely to result in improved incidence surveillance over methods currently in wide use. Reasonably accurate and precise age-specific estimates of incidence are important to target better prevention, diagnosis and care strategies.

**Fig. 4.**
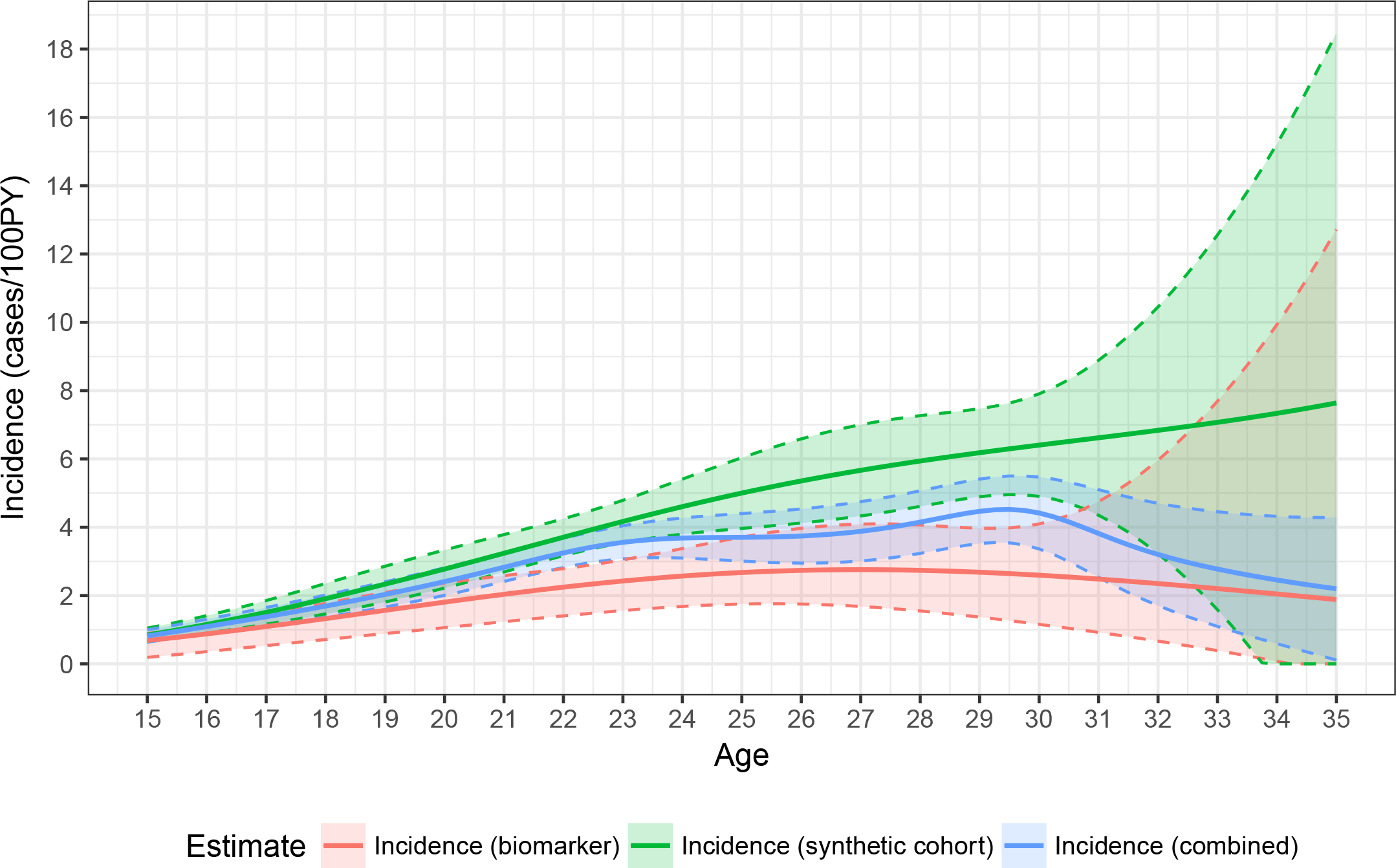
HIV incidence by age in males and females aged 15-35, using the synthetic cohort, recency biomarker and combined methods.

## Acknowledgments

The authors gratefully acknowledge the Centre for High Performance Computing (CHPC), South Africa, for providing computational resources to this research project.

The authors are grateful to the Consortium for the Evaluation and Performance of HIV Incidence Assays (CEPHIA) for allowing us to use its recent infection biomarker calibration data in this study, specifically the principal investigators Gary Murphy, Christopher D. Pilcher, Michael P. Busch and Alex Welte, and the core team comprising Sheila Keating, Mila Lebedeva, Dylan Hampton, Jake Hall, Elaine McKinney, Kara Marson, Shelley Facente, Eduard Grebe, Reshma Kassanjee and Trust Chibawara.

CEPHIA comprises: Oliver Laeyendecker, Thomas Quinn, David Burns (National Institutes of Health); Alex Welte, Eduard Grebe, Reshma Kassanjee, David Matten, Hilmarie Brand, Trust Chibawara (South African Centre for Epidemiological Modelling and Analysis); Gary Murphy, Elaine Mckinney, Jake Hall (Public Health England); Michael Busch, Sheila Keating, Mila Lebedeva, Dylan Hampton (Blood Systems Research Institute); Christopher Pilcher, Kara Marson, Shelley Facente, Jeffrey Martin; (University of California, San Francisco); Susan Little (University of California, San Diego); Anita Sands (World Health Organization); Tim Hallett (Imperial College London); Sherry Michele Owen, Bharat Parekh, Connie Sexton (Centers for Disease Control and Prevention); Matthew Price, Anatoli Kamali (International AIDS Vaccine Initiative); Lisa Loeb (The Options Study – University of California, San Francisco); Jeffrey Martin, Steven G Deeks, Rebecca Hoh (The SCOPE Study – University of California, San Francisco); Zelinda Bartolomei, Natalia Cerqueira (The AMPLIAR Cohort – University of São Paulo); Breno Santos, Kellin Zabtoski, Rita de Cassia Alves Lira (The AMPLIAR Cohort – Grupo Hospital Conceição); Rosa Dea Sperhacke, Leonardo R Motta, Machline Paganella (The AMPLIAR Cohort – Universidade Caxias Do Sul); Esper Kallas, Helena Tomiyama, Claudia Tomiyama, Priscilla Costa, Maria A Nunes, Gisele Reis, Mariana M Sauer, Natalia Cerqueira, Zelinda Nakagawa, Lilian Ferrari, Ana P Amaral, Karine Milani (The São Paulo Cohort – University of São Paulo, Brazil); Salim S Abdool Karim, Quarraisha Abdool Karim, Thumbi Ndungu, Nelisile Majola, Natasha Samsunder (CAPRISA, University of Kwazulu-Natal); Denise Naniche (The GAMA Study – Barcelona Centre for International Health Research); Inácio Mandomando, Eusebio V Macete (The GAMA Study – Fundacao Manhica); Jorge Sanchez, Javier Lama (SABES Cohort – Asociación Civil Impacta Salud y Educacion (IMPACTA)); Ann Duerr (The Fred Hutchinson Cancer Research Center); Maria R Capobianchi (National Institute for Infectious Diseases “L. Spallanzani”, Rome); Barbara Suligoi (Istituto Superiore di Sanità, Rome); Susan Stramer (American Red Cross); Phillip Williamson (Creative Testing Solutions / Blood Systems Research Institute); Marion Vermeulen (South African National Blood Service); and Ester Sabino (Hemocentro do Sao Paolo).

## Appendix 1: Optimal RITA identification and calibration

The data used in the calibration and contextual adaptation of the Recent Infection Testing Algorithm (RITA) used in this study were provided by the CEPHIA Group. The UCSF Human Research Protection Program and IRB (formerly CHR, #10-02365) approved the study procedures.

The performance of a test for recent infection or RITA for the purposes of incidence estimation is captured in two parameters: The Mean Duration of Recent Infection (MDRI) and the False-Recent Rate (FRR).

The *MDRI* is the average amount of time that individuals spend exhibiting the ‘recent’ biomarker, while infected for less than some cut-off time (denoted *T*, 2 years in the present work). This captures the defining biological aspects of the recency test. The *FRR* is the proportion of individuals infected for longer than the cut-off time *T*, but who nevertheless produce a recent result on the test. FRR is inevitably context-dependent, and critically depends on epidemiological factors such as the prevalence of HIV infection and antiretroviral treatment coverage. The latter is important, because with the reduction in antigenic pressure associated with viral suppression, immune markers tend to revert to a state similar to early infection, resulting in ‘false’ recent classifications. Inclusion of a viral load threshold in a RITA reduces the FRR in treated individuals to close to zero.

In order to estimate MDRI and context-specific FRR for a given RITA, the chosen recent infection case definition was applied to LAg, Bio-Rad Avidity and viral load results on the CEPHIA evaluation panel. The CEPHIA evaluation panel consists of 2,500 well-characterised specimens and was employed in the independent evaluation of a range of candidate tests for recent infection, including both the LAg and Bio-Rad Avidity assays. In order to define an ‘optimal’ RITA, a range of combinations of thresholds on three available biomarkers – the LAg normalised optical density (ODn), the Bio-Rad Avidity index (AI) and the viral load (copies/ml) – were applied and MDRI and context-dependent FRR estimated for the epidemiological context of this study.

MDRI was estimated by fitting a regression model for the probability of testing recent as a function of estimated time since infection, *P*_*R*_(*t*), using a logit link function and a cubic polynomial in time (since estimated date of detectable infection), to CEPHIA evaluation panel data. The function was fit to data points up to 800 days post-infection. The MDRI was then obtained by integrating the function from 0 to *T*. Confidence intervals were obtained by resampling subjects in 10,000 bootstrap iterations. MDRI was estimated on subtype C-infected specimens only, the predominant subtype in the surveyed population. The estimated MDRI was adjusted for the sensitivity of the screening algorithm used in this study, namely NAAT in pools of five specimens (effective detection threshold of 500 copies/ml), i.e. 7.7 days shorter MDRI than estimated using the CEPHIA reference test [27].

Estimating *context-dependent FRR* requires defining the epidemiological context, namely HIV prevalence, HIV incidence and treatment coverage and then estimating FRR in untreated and treated individuals separately, combining the estimates into a weighted average according to treatment coverage in the population. To obtain contextual epidemiological parameters, the survey data was analysed to obtain prevalence and treatment coverage proportions, and an initial incidence analysis was conducted, using a standard RITA comprising LAg (≤ 1.5) and viral load (> 100 copies/ml), with MDRI and a crude FRR estimated from CEPHIA data, to obtain an overall incidence in the population of interest. ARV testing was not included in the RITA, as this may have an unknown impact on MDRI, and earlier work using this survey demonstrated limited benefit for FRR and precision of incidence estimates [21].

These parameters were then employed to estimate context-specific FRR, by estimating the FRR in untreated individuals and treated individuals separately, and weighting these estimates according to treatment coverage. Confidence intervals were obtained by resampling subjects in 20,000 bootstrap iterations. For untreated individuals, the function *P*_*R*_(*t*) was fit using CEPHIA data for subtype C-infected individuals, from all times post-infection, and weighted according to the probability density function for times since infection in the untreated population. The distribution of times since infection was parameterised as a Weibull survival function (i.e. remaining in the untreated state), with the shape and scale parameters chosen to produce the desired treatment coverage in a population with the specified incidence and prevalence and scaled to recent incidence. The FRR in treated subjects, *P*_*R*|*tx*_ is simply the binomially estimated probability that treated subjects infected for longer than *T* would produce a recent result,^1^ since the FRR in treated subjects appears not to depend strongly on time since infection. The weighted FRR estimate was obtained as shown in Eq 15:

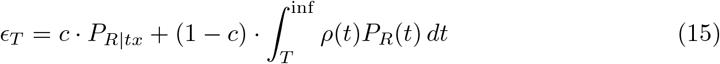

where *c* is the treatment coverage, 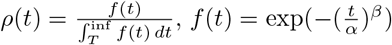 and *α* and *β* the Weibull scale and shape parameters, respectively. This approach was previously described in [28].

We obtained incidence estimates for a range of LAg and Bio-Rad Avidity threshold combinations, together with a viral load threshold of 75 copies/ml, and evaluated relative standard error (RSE) on the incidence estimate. Reproducibility of the incidence estimate was obtained from 100,000 bootstrap iterations, each drawing from the distributions of test property estimates, HIV prevalence and prevalence of recency among HIV-positives (from the survey dataset, analysed using the survey R package [29] to account for the complex sampling frame), using the inctools R package [23]. We adopted the ‘optimal’ recency case definition of: (NAAT-positive and antibody negative) OR (antibody positive, LAg ODn ≤ 2.5, Bio-Rad Avidity AI ≤ 30 and viral load > 75 copies/ml). A selection of threshold combinations is shown in Table 5.

**Table 4.** Precision of incidence estimate produced by a range of LAg, Bio-Rad Avidity and viral load threshold combinations.

| LAg ODn<br>≤ | Bio-Rad AI<br>≤ | Viral Load<br>$>$ | MDRI*<br>(95% CI)<br>*days* | Context-adapted FRR<br>(95% CI)<br>*%* | RSE on incidence<br>*%* |
| --- | --- | --- | --- | --- | --- |
| 1.5 | 20 | 75 | 156 (136,179) | 0.06% (0.00%,0.17%) | 18.90% |
| 1.5 | 30 | 75 | 172 (148,197) | 0.06% (0.00%,0.17%) | 18.70% |
| 1.5 | 40 | 75 | 176 (152,201) | 0.12% (0.01%,0.31%) | 19.20% |
| 1.5 | 50 | 75 | 179 (155,205) | 0.12% (0.01%,0.31%) | 19.20% |
| 1.5 | 60 | 75 | 179 (155,205) | 0.13% (0.01%,0.33%) | 19.40% |
| 1.75 | 20 | 75 | 161 (140,183) | 0.09% (0.02%,0.22%) | 18.20% |
| 1.75 | 30 | 75 | 178 (155,204) | 0.09% (0.02%,0.22%) | 18.00% |
| 1.75 | 40 | 75 | 186 (160,215) | 0.16% (0.02%,0.39%) | 18.70% |
| 1.75 | 50 | 75 | 190 (164,219) | 0.16% (0.02%,0.40%) | 18.70% |
| 1.75 | 60 | 75 | 193 (166,223) | 0.17% (0.03%,0.41%) | 19.10% |
| 2 | 20 | 75 | 172 (149,196) | 0.09% (0.02%,0.22%) | 18.00% |
| 2 | 30 | 75 | 191 (166,218) | 0.09% (0.02%,0.22%) | 17.80% |
| 2 | 40 | 75 | 200 (174,229) | 0.16% (0.02%,0.39%) | 18.40% |
| 2 | 50 | 75 | 205 (177,235) | 0.16% (0.02%,0.39%) | 18.50% |
| 2 | 60 | 75 | 208 (179,238) | 0.18% (0.02%,0.41%) | 18.70% |
| 2.25 | 20 | 75 | 186 (164,210) | 0.11% (0.03%,0.25%) | 17.20% |
| 2.25 | 30 | 75 | 210 (184,236) | 0.12% (0.03%,0.27%) | 17.10% |
| 2.25 | 40 | 75 | 222 (192,254) | 0.21% (0.05%,0.47%) | 18.20% |
| 2.25 | 50 | 75 | 232 (201,265) | 0.22% (0.05%,0.47%) | 18.10% |
| 2.25 | 60 | 75 | 237 (206,269) | 0.42% (0.08%,1.14%) | 20.50% |
| 2.5 | 20 | 75 | 194 (171,219) | 0.14% (0.04%,0.31%) | 16.80% |
| **2.5** | **30** | **75** | **217 (192,244)** | **0.17% (0.05%,0.35%)** | **16.60%** |
| 2.5 | 40 | 75 | 233 (204,265) | 0.30% (0.09%,0.61%) | 17.90% |
| 2.5 | 50 | 75 | 249 (218,282) | 0.30% (0.10%,0.61%) | 17.60% |
| 2.5 | 60 | 75 | 260 (227, 293) | 0.51% (0.15%,1.26%) | 19.30% |

\*MDRI adjusted for sensitivity of screening algorithm

## Appendix 2: Sensitivity analyses

## Sensitivity of incidence estimates to False-Recent Rate

To evaluate the sensitivity of incidence estimates to the false-recent rate (FRR), we repeated the estimation procedure using values of FRR ranging from 0% to 1%. With higher FRR values, the biomarker-based estimates of incidence are lower, and this impact is greatest at the higher ages (which also have higher prevalence). Fig 5 shows the age-continuous biomarker-based estimates for demonstrative ages (ages 15, 20, 25 and 30) for both males and females. For this analysis a relative standard error (RSE) on the FRR estimate of 50% was assumed (similar to the estimated value used in the primary analysis).

Estimates using the combined method for FRR values ranging from 0% to 1% and RSE on FRR of 50% are shown in Fig 6. In these plots the incidence estimates counter-intuitively do not decline with higher FRRs. This results from greater uncertainty in the biomarker-based estimates, owing to the large RSE on FRR, which results in the combined estimates being weighted towards the synthetic cohort-based estimates (which are higher). To avoid this artefact, the analysis was repeated assuming an RSE on FRR of 0%, shown in Fig 7.

**Fig. 5.**
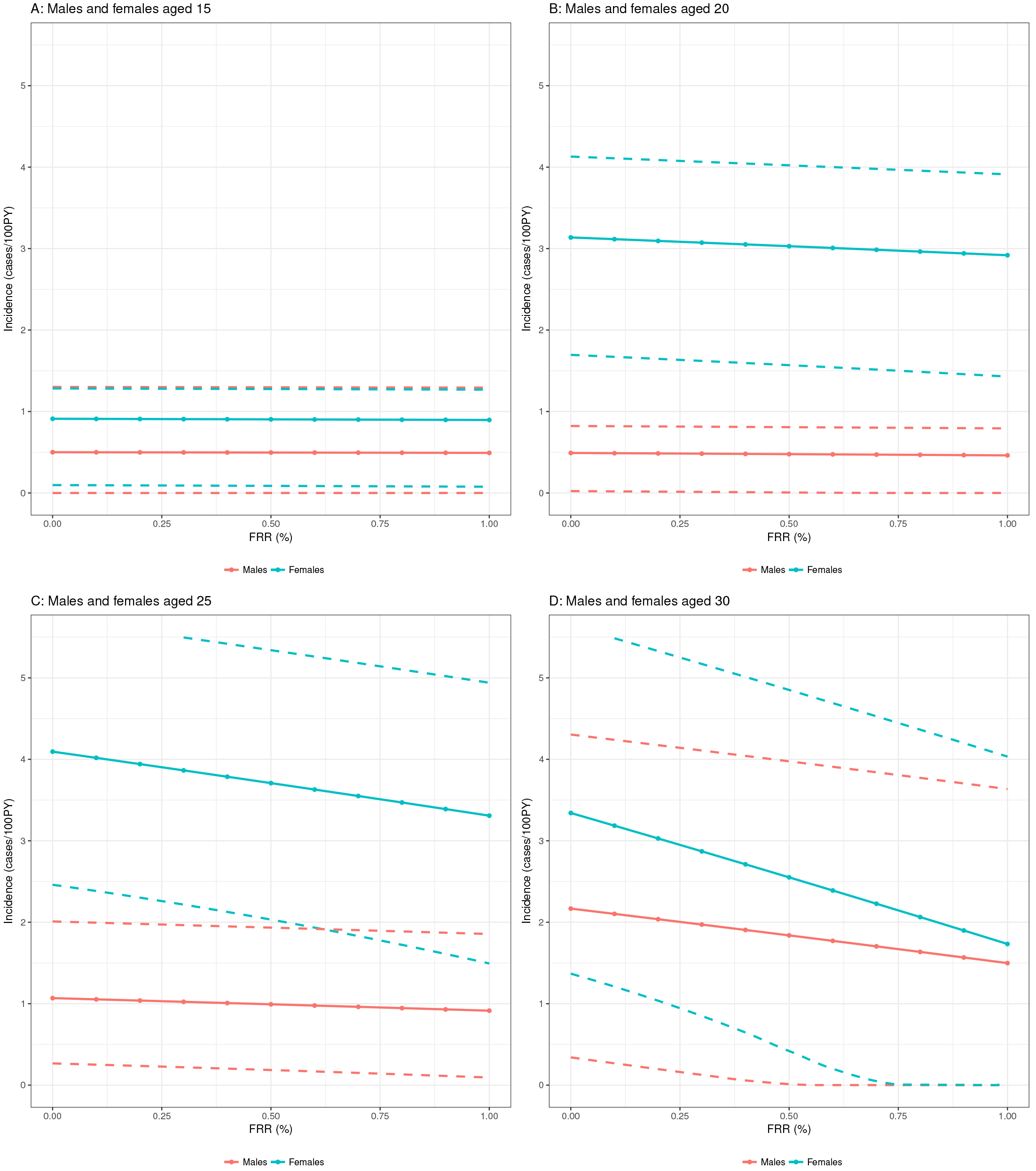
Incidence estimates (biomarker method) for a range of FRR values (relative standard error on FRR of 50%).

**Fig. 6.**
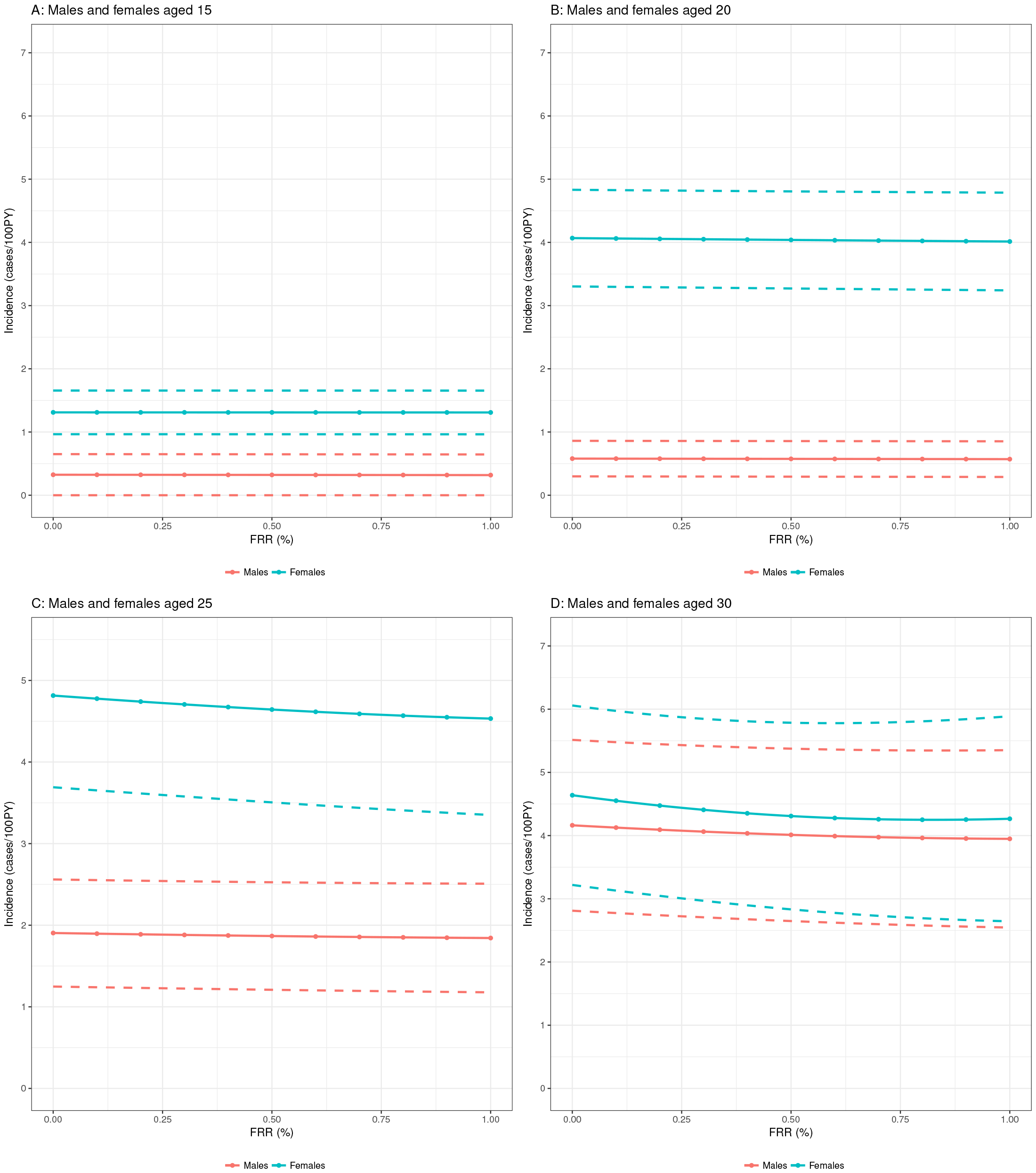
Incidence estimates (combined method) for a range of FRR values (relative standard error on FRR of 50%).

**Fig. 7.**
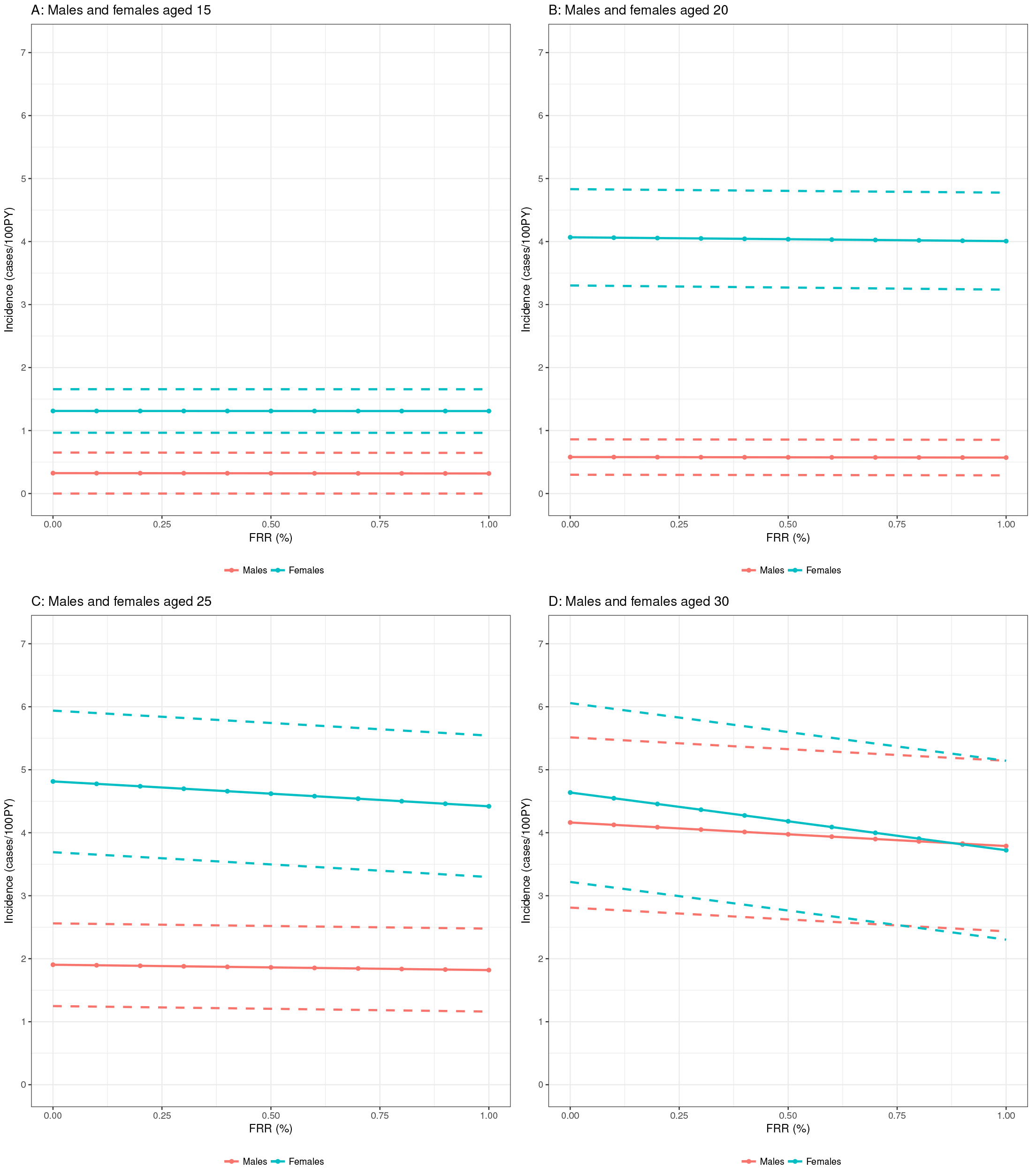
Incidence estimates (combined method) for a range of FRR values (relative standard error on FRR of 0%).

## Sensitivity of average incidence estimates to weighting scheme

Calculating ‘average incidence’ over a particular age range requires estimating sex and age-specific incidence and weighting the estimates over the age range by either the sampling density or the susceptible population density (see Methods section). In Table 5, we report the average incidence obtained for a set of age ranges when weighting by the sampling or susceptible population densities. Only the susceptible population-weighted estimates are reported in the main text of the paper.

**Table 5.** Average incidence estimates (combined method), weighted by susceptible population density and sampling density.

| Age group<br>*years* | Weighted by susceptible population density |  |  | Weighted by sampling density |  |  |
| --- | --- | --- | --- | --- | --- | --- |
|  | Males<br>P.E. (95% CI)<br>*cases/100PY* | Females<br>P.E. (95% CI)<br>*cases/100PY* | Total<br>P.E. (95% CI)<br>*cases/100PY* | Males<br>P.E. (95% CI)<br>*cases/100PY* | Females<br>P.E. (95% CI)<br>*cases/100PY* | Total<br>P.E. (95% CI)<br>*cases/100PY* |
| [15,20) | 0.36 (0.00,0.61) | 2.62 (2.07,3.11) | 1.51 (1.13,1.75) | 0.37 (0.04,0.61) | 2.77 (2.17,3.28) | 1.46 (1.09,1.69) |
| [20,25) | 1.22 (0.71,1.49) | 4.73 (4.00,5.54) | 2.95 (2.46,3.28) | 1.16 (0.67,1.42) | 4.72 (3.98,5.51) | 2.94 (2.46,3.27) |
| [25,30) | 2.64 (1.13,3.26) | 4.65 (3.68,5.79) | 3.61 (2.48,4.19) | 2.72 (1.16,3.36) | 4.65 (3.67,5.81) | 3.79 (2.65,4.37) |
| [30,35) | 2.76 (0.38,4.7+9) | 3.56 (1.77,6.15) | 3.14 (1.38,4.65) | 2.88 (0.48,4.64) | 3.62 (1.89,6.11) | 3.29 (1.54,4.73) |
| [15,30) | 1.26 (0.64,1.49) | 3.83 (3.35,4.37) | 2.54 (2.07,2.77) | 1.14 (0.55,1.36) | 3.94 (3.41,4.51) | 2.52 (2.09,2.77) |

As can be seen in Table 5, average incidence estimates for 5-year age bins, as well as the entire 15-30 age group are not very sensitive to weighting scheme. In all cases the point estimate for average incidence weighted by sampling density falls within the 95% confidence interval of the susceptible population density-weighted estimates, and the differences in point estimates are very small.

## Sensitivity of incidence estimates to time-varying age structure of prevalence

We have no direct information from the cross-sectional survey on the prevalence gradient in time (i.e. the partial derivative of prevalence with respect to time), as required by the original Mahiance et al. synthetic cohort incidence estimator:

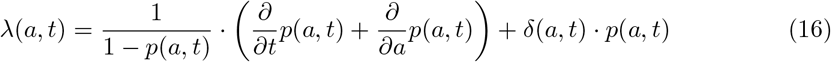

We therefore repeated the entire estimation procedure, assuming various age-specific prevalence gradients in time, 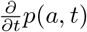, at the time of the survey. The THEMBISA model‘s age-specific prevalence estimates for the period 2010 to 2014 indicate a stable epidemic in persons aged 15-35, with approximately linear change and gradients varying between −0.009 and 0.010 in females and between −0.013 and 0.004 in males in the various one-year age bins.

We parameterised assumed change in age-specific prevalence by age as an exponential decline (or increase) in the form *p*(*a*, *t*) = *p*(*a*) · *e*^−γ*t*^ so that 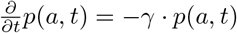. A steep decline in prevalence (halving of prevalence in approximately 10 years) is obtained with γ = 0.07. We repeated the analysis for males, females and overall (using data for all ages, but reporting results for age-specific incidence at ages 15, 20, 25 and 30) using four values, representing rapidly increasing and decreasing age-specific prevalence and more plausible time-gradients of prevalence, as well as the extreme case where prevalence is plummeting (with a half-life of approximately 5 years, γ = 0.14):

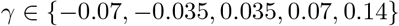

The results, compared to the primary analysis, assuming 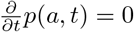, are shown in Table 6.

**Table 6.**
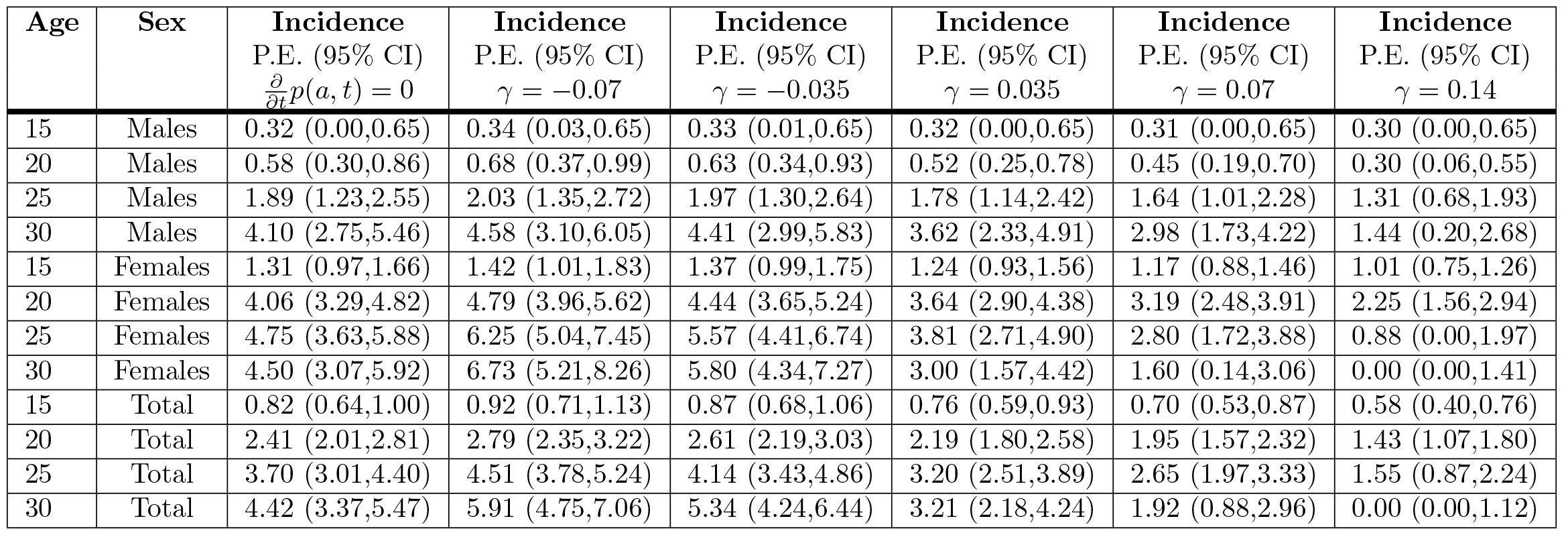
Average incidence estimates (combined method), weighted by susceptible population density and sampling density.

| Age | Sex | Incidence<br>P.E. (95% CI)<br>$\frac{\partial}{\partial t}p(a, t) = 0$ | Incidence<br>P.E. (95% CI)<br>$\gamma = -0.07$ | Incidence<br>P.E. (95% CI)<br>$\gamma = -0.035$ | Incidence<br>P.E. (95% CI)<br>$\gamma = 0.035$ | Incidence<br>P.E. (95% CI)<br>$\gamma = 0.07$ | Incidence<br>P.E. (95% CI)<br>$\gamma = 0.14$ |
| --- | --- | --- | --- | --- | --- | --- | --- |
| 15 | Males | 0.32 (0.00,0.65) | 0.34 (0.03,0.65) | 0.33 (0.01,0.65) | 0.32 (0.00,0.65) | 0.31 (0.00,0.65) | 0.30 (0.00,0.65) |
| 20 | Males | 0.58 (0.30,0.86) | 0.68 (0.37,0.99) | 0.63 (0.34,0.93) | 0.52 (0.25,0.78) | 0.45 (0.19,0.70) | 0.30 (0.06,0.55) |
| 25 | Males | 1.89 (1.23,2.55) | 2.03 (1.35,2.72) | 1.97 (1.30,2.64) | 1.78 (1.14,2.42) | 1.64 (1.01,2.28) | 1.31 (0.68,1.93) |
| 30 | Males | 4.10 (2.75,5.46) | 4.58 (3.10,6.05) | 4.41 (2.99,5.83) | 3.62 (2.33,4.91) | 2.98 (1.73,4.22) | 1.44 (0.20,2.68) |
| 15 | Females | 1.31 (0.97,1.66) | 1.42 (1.01,1.83) | 1.37 (0.99,1.75) | 1.24 (0.93,1.56) | 1.17 (0.88,1.46) | 1.01 (0.75,1.26) |
| 20 | Females | 4.06 (3.29,4.82) | 4.79 (3.96,5.62) | 4.44 (3.65,5.24) | 3.64 (2.90,4.38) | 3.19 (2.48,3.91) | 2.25 (1.56,2.94) |
| 25 | Females | 4.75 (3.63,5.88) | 6.25 (5.04,7.45) | 5.57 (4.41,6.74) | 3.81 (2.71,4.90) | 2.80 (1.72,3.88) | 0.88 (0.00,1.97) |
| 30 | Females | 4.50 (3.07,5.92) | 6.73 (5.21,8.26) | 5.80 (4.34,7.27) | 3.00 (1.57,4.42) | 1.60 (0.14,3.06) | 0.00 (0.00,1.41) |
| 15 | Total | 0.82 (0.64,1.00) | 0.92 (0.71,1.13) | 0.87 (0.68,1.06) | 0.76 (0.59,0.93) | 0.70 (0.53,0.87) | 0.58 (0.40,0.76) |
| 20 | Total | 2.41 (2.01,2.81) | 2.79 (2.35,3.22) | 2.61 (2.19,3.03) | 2.19 (1.80,2.58) | 1.95 (1.57,2.32) | 1.43 (1.07,1.80) |
| 25 | Total | 3.70 (3.01,4.40) | 4.51 (3.78,5.24) | 4.14 (3.43,4.86) | 3.20 (2.51,3.89) | 2.65 (1.97,3.33) | 1.55 (0.87,2.24) |
| 30 | Total | 4.42 (3.37,5.47) | 5.91 (4.75,7.06) | 5.34 (4.24,6.44) | 3.21 (2.18,4.24) | 1.92 (0.88,2.96) | 0.00 (0.00,1.12) |

Estimates are not very sensitive to plausible time-gradients of age-specific prevalence at ages where the prevalence is relatively low. The impact – as would be expected given the assumption of exponential increase or decline – is substantial at ages with higher estimated prevalence, especially females over 20 years of age. Model-based estimates of age-specific prevalence suggest modest and linear declines at most ages in the range of interest, with a small positive gradient at very young ages (likely reflecting the ageing-in of perinatally-infected individuals, and therefore not likely to reflect higher incidence in this population) and small negative gradient at the ages with the highest incidence, indicating that incidence may be very slightly over-estimated if such a decline is taking place in the surveyed population.

## Footnotes

1 In the CEPHIA Evaluation Panel, all treated subjects were virally suppressed, resulting in an estimate of *P*_*R*|*tx*_ = 0 in all cases where a supplemental viral load threshold is applied. In real-world populations, it is likely that a certain (unknown) proportion of treated subjects would be virally unsuppressed and that the FRR in treated subjects would therefore be non-zero.

